# A novel glia-immune humanized mouse model for investigating neuroimmune mechanisms in CNS HIV infection and diverse neurological disorders

**DOI:** 10.1101/2025.03.05.641678

**Authors:** Shallu Tomer, Lin Pan, Jeffrey Harding, Valerie Rezek, Mitchell Krawczyk, Ethan Cook, Nandita Kedia, Li Wang, Jonathan Le, Christopher Platt, Heather Martin, Erik Gramajo, Xiang Li, Yewen Zhang, Ariel S. Peng, Yi Lu, Wenli Mu, Jing Wen, Scott Kitchen, Ye Zhang, Anjie Zhen

**Author notes:** Co-first authors (Shallu Tomer, Lin Pan). Corresponding Authors: Anjie Zhen, Ye Zhang.

## Abstract

Dysregulation in neuroimmune interactions drives pathology in diverse neurological disorders. However, human cell-based in vivo models are limited for clinically relevant mechanistic studies and drug testing. To address this, we developed a novel humanized mouse model integrating all four major types of human glia - astrocytes, oligodendrocyte precursor cells (OPCs), oligodendrocytes, and microglia - in the brain, alongside donor-matched human immune system in peripheral blood and lymphoid tissues. We applied this model to study HIV-associated neurocognitive disorders, which can persist in tissue reservoirs including in the brain despite viral suppression by antiretroviral therapy. These glia-immune humanized mice supported robust HIV-1 replication in the peripheral blood, lymphoid tissues, and the brain, and recapitulated the synapse loss observed in patients. Furthermore, HIV-infected mice exhibited heightened inflammation in both peripheral and brain tissues. Notably, OPCs and oligodendrocytes in HIV-infected brains adopted an immune activated phenotype, upregulating interferon-stimulated genes, highlighting understudied roles of oligodendroglia in HIV infection. RNA-sequencing of human glia identified the upregulation of inflammasome-associated genes and the downregulation of genes involved in transcriptional, epigenetic, and metabolic regulation. Overall, this model provides a powerful in vivo platform to investigate human neuroimmune interactions relevant to diverse neurological disorders, offering critical insights into HIV CNS infection, pathology, and potential therapeutic strategies targeting CNS reservoirs and neuroinflammatory pathways.

## Introduction

Neuroinflammation, glial dysfunction, and immune dysregulation are key drivers of disease progression in a variety of neurological disorders, including Alzheimer’s disease (AD), Parkinson’s disease (PD), Huntington’s disease (HD), stroke, and HIV-associated neurocognitive disorder (HAND)^1,2^. Despite their critical roles and potential as therapeutic targets, investigation of neuroimmune dysfunction is among the most challenging research areas, largely due to the lack of clinically relevant *in vivo* models. Human and mouse glia are more distinct in transcriptome than their neuronal counterparts ^3,4^. They react differently to key pathological features of neurological diseases, such as exacerbated neuroinflammation and oxidative stress^3^. Similarly, human and mouse immune cells diverge in significant ways. Consequently, conventional mouse models do not capture human-specific features of the neuro-immune systems. The vast majority of drugs identified in mouse models fail in clinical trials, severely delaying progress. While human iPSC-derived in vitro cultures are useful for investigating one or several types of human cells, they lack the full collection of glia and immune cell types and do not reproduce key anatomical structures such as the blood-brain barrier. Humanized mouse models incorporating specific human immune and glial cells have shown promise for investigating the functions and dysfunctions of these human cell types *in vivo* ^5–9^. However, none of the current humanized mouse models integrate all major human immune and glial cell types. This represents a critical limitation, as resident glial cells and infiltrating immune cells continuously interact and mutually influence each other’s behavior, with these interactions being essential determinants of neuroinflammation outcomes, disease progression, and brain repair. Thus, the development of a humanized mouse model encompassing HLA-matched full complement of human glial and immune cell types would provide an indispensable platform for mechanistic studies of human neuroimmune interactions in vivo and for preclinical testing of therapeutics across a range of neurological disorders.

Despite development of effective combined anti-retroviral therapy (ART), HIV remains largely incurable due to persistent viral reservoirs, including in the central nervous system (CNS). As many as 50% of people living with HIV (PLWH) experience CNS comorbidities ^10^. Compared to peripheral reservoirs, our understandings of the infection and reservoirs in the CNS are limited. Pathogenesis of HIV infection in the CNS is multifactorial: HIV replicates and persists in the CNS resident cells, viral proteins are neurotoxic, CNS inflammation occurs and many other pathogenic effects are observed. While microglia appear to be the main HIV-1 reservoir in the brain^11^, astrocytes and oligodendroglia - including oligodendrocytes and oligodendrocyte precursor cells (OPCs) - may also be affected^12–14^. Astrocytes are essential for brain homeostasis and can drive the induction and progression of a pro-inflammatory environment. Damage to oligodendroglia and myelin has been detected in patients with HIV, correlating with neurocognitive deficits ^14,15^. Therefore, understanding how infiltrating immune cells and resident CNS glia - microglia, astrocytes, and oligodendroglia - support and respond to HIV infection is critical for elucidating the pathogenesis of CNS comorbidity.

Commonly used platforms for studying HIV CNS infection and pathogenesis have limitations. HIV does not productively infect mice or non-human primates (NHPs)^16^, and obtaining HIV-infected human brain tissues from people living with HIV (PLWH) is challenging. As a result, the molecular mechanisms of HIV CNS infection and CNS pathogenesis remain largely elusive. Humanized mice transplanted with human hematopoietic stem cells (HSCs) allow the development of a functional human immune system, support robust HIV replication, and enable HIV latency establishment^17–22^. These models have proven valuable for investigating HIV immunopathogenesis, latency reactivation and testing gene and cell therapies as described in ours^17–19,23–28^ and others’ studies^17,19,20,27–33^. However, available models, including new humanized mouse models that contain injected astrocyte^34^, or express human IL-34 to support better development of microglia-like cells^34–37^, do not include all four types of human CNS glia (microglia, astrocytes, OPCs, and oligodendrocytes) and donor-matched human immune cells.

While much of HAND research has traditionally focused on neurons, and more recently on microglia and astrocytes, oligodendroglia have been largely overlooked. In PLWH, abnormalities in white matter, where oligodendrocytes and myelin are enriched, is a prominent neuropathological feature observed in both pre- and post-ART eras and is associated with cognitive impairment^38^. Viral proteins such as Tat and gp120 and neuroinflammation impair oligodendrocyte maturation and survival, while HIV-induced neuroinflammation exacerbates demyelination and inhibits remyelination^13,14^. Recent scRNAseq studies of postmortem human brain tissues from ART-treated PLWH also revealed significant transcriptomic changes in OPCs ^39^. However, previous mechanistic studies of CNS HIV infection have primarily relied on in vitro cell cultures and transgenic mouse models that overexpress individual HIV proteins, such as Tat or gp120 ^40,41^. It is challenging to capture the full dynamics of viral replication, latency, and immune interactions seen in human infection using these approaches, as they focus on isolated proteins at artificially high levels - more akin to severe HIV encephalitis, which has become increasingly rare in the ART era. Consequently, little is known about how HIV impacts OPC and oligodendrocyte function in vivo.

To address these critical knowledge gaps, we applied our novel glia-immune humanized mice to investigate HIV CNS comorbidities. These glia-immune humanized mice support robust HIV infection in both the periphery and the CNS. Chronic immune activation and neuroinflammation are strongly associated with increased incidence of HAND^42^. Consistent with this, we observed elevated inflammation and significant transcriptomic changes in human glia cells within the brains of infected mice, including elevated type I interferon (IFN-I) responses, which mimic human conditions. Notably, OPCs and oligodendrocytes in HIV-infected brains adopted an immune activated phenotype, upregulating interferon-stimulated genes, highlighting understudied roles of oligodendroglia in HIV infection. Our model provides a unique platform to investigate the interplay between HIV infection, chronic immune activation, and neuroinflammation within a physiologically relevant humanized CNS environment. Beyond HAND, this novel humanized mouse model will be an indispensable tool for mechanistic investigation of neuroimmune interactions and preclinical therapeutic testing in a broad spectrum of neurological disorders, including AD, PD, HD, stroke, and brain tumors.

## Results

### The glia-immune humanized NSG mouse model supports the development of human multilineage immune cells

To study neuroimmune interactions in HIV CNS infection and other diseases and the role of human glia in CNS inflammation, we developed a novel glia-immune humanized mouse model. In this model, mice are reconstituted with donor-matched human glial cells in the brain and a human immune system in periphery and lymphoid tissues. As shown in **Fig. 1A**, neonatal immunodeficient NSG mice (NOD.Cg-*Prkdc^scid^Il2rg^tm1Wjl^* /SzJ) were injected with donor-matched gestation week <18 liver-derived CD34+ HSCs (via intrahepatic injection) and brain-derived astrocytes and OPCs (via intracranial injection, as detailed in^3^). For glial transplantation, astrocytes and OPCs were injected into the neonatal mouse brain at 4 injection sites, including the striatum and thalamus bilaterally. Mice were bled and the reconstitution level was assessed 14-18 weeks after transplant.

**Figure 1.**
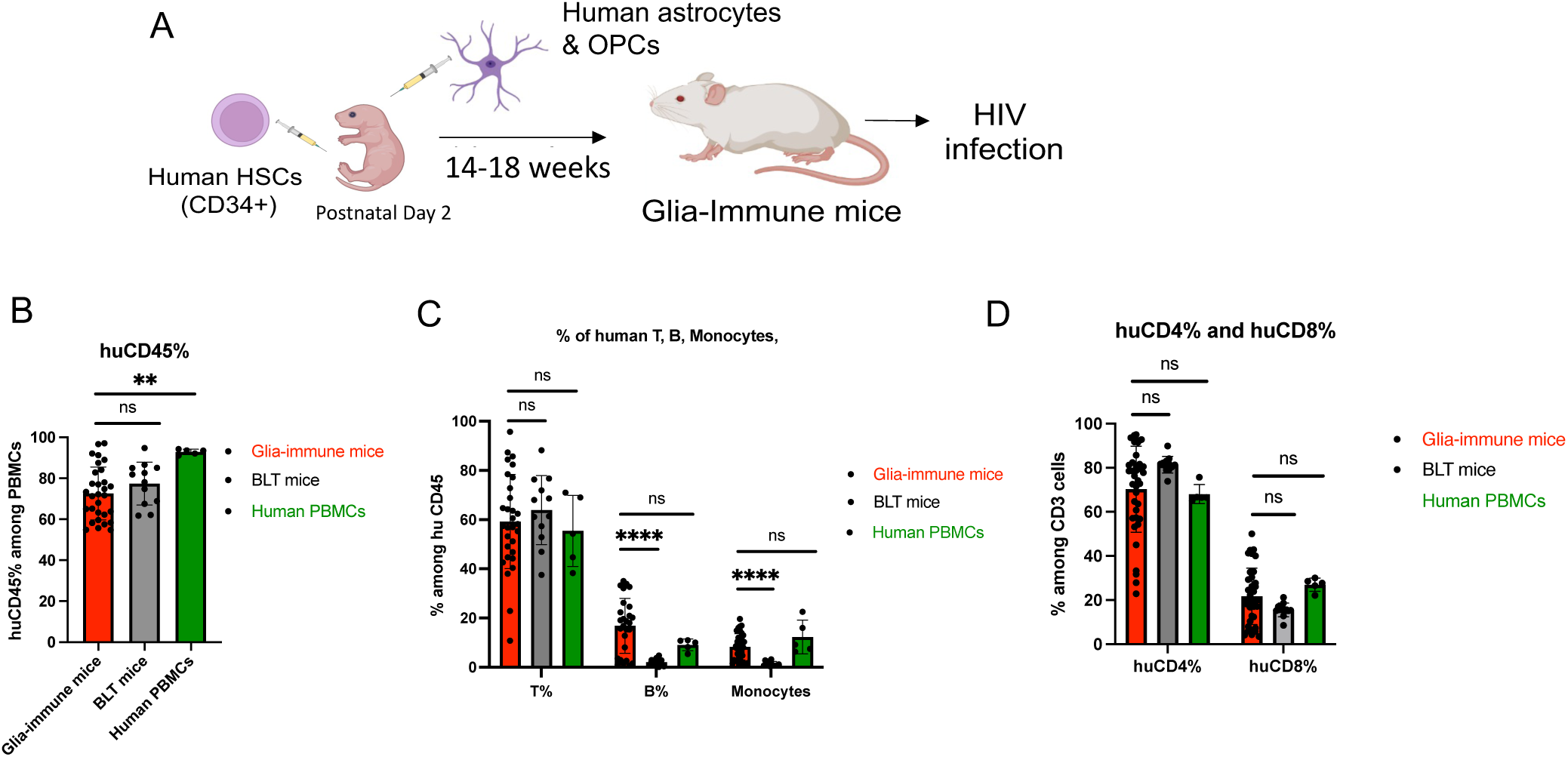
Successful engraftment and differentiation of human immune and glial cells in glia-immune humanized mice. A) Human astrocytes and OPCs were transplanted into the subcortical areas of the brains at 1-2mm depth and donor matched human CD34+ HSCs were co-transplanted into the livers of immunodeficient mice at postnatal Day 2. 14-18 weeks after reconstitution, mice were bled and human immune reconstitution was assessed by flow-cytometry by staining with human antibodies. B) Percentage of huCD45+ cells in PBMCs from the glia-immune humanized mice (n=30), humanized BLT mice (n=12) or primary human PBMCs from healthy donors (n=5). C) Percentage of human T cells (huCD3), B cells (huCD19), Monocytes (huCD14) among huCD45 positive cells. D) Percentage of human T-Helper cells (huCD4) and T-cytotoxic cells (huCD8) among T cells. **p=0.0056, ****p<0.0001 by Mann-Whitney test.

We first determined whether glia-immune humanized mice exhibit similar levels of human immune reconstitution when compared to primary human peripheral blood mononuclear cells (PBMCs) from healthy donors and the Bone marrow-Liver-Thymus (BLT) humanized mouse, a well characterized humanized mouse model that supports robust immune reconstitution and allows for comprehensive studies in HIV immunity. The BLT humanized mouse model is key for seminal studies of cell and gene therapy for HIV cure^8,18,20,43–54^, studies in HIV latency^21,22,55,56^ and mechanistic studies of HIV immunopathogenesis ^27,51,52,57–61^. We performed a comparative analysis among humanized glia-immune mice, BLT mice, and fresh human PBMCs of the percentage of human leukocytes (CD45+ cells) in the PBMCs and the distribution of human immune cell subsets, including T cells (CD3+), B cells (CD19+), monocytes (CD14+) using flow cytometry.

In the glia-immune humanized mice, we observed similar level of CD45+% cells among the PBMCs as compared to BLT mice **(Fig. 1B).** Within the human CD45+ leukocytes of the glia-immune mice, we observed successful development of T cells (CD3+), B cells (CD19+) and monocytes (CD14+), at levels similar to primary human PBMCs from healthy donors **(Fig 1C).** Glia-immune humanized mice showed significantly better development of B cells and monocytes compared to BLT mice **(Fig 1C).** Among the T cell subsets, 70.3% of T cells in the glia-immune humanized mice were CD4+ and 21.7% were CD8+, similar to primary human PBMCs and to BLT mice **(Fig 1D).** These results suggest that glia-immune humanized mice support robust development of multiple hematopoietic lineages, resembling human PBMCs more closely than the humanized BLT mice.

### Reconstitution of human microglia-like cells, astrocytes, OPCs, and oligodendrocytes

To examine the reconstitution of human glia in host mouse brain, we performed IHC analyses of xenografted mouse brains at 26 weeks post reconstitution. As shown in **Fig. 2A**, we observed widespread engraftment of human glia in the brains, including astrocytes (hNuclei+Sox9+) (**Fig. 2D**), microglia-like cells (huNuclei+Iba1+) (**Fig. 2E**), Oligodendroglia (huNuclei+Olig2+, **Fig 2F**), including OPCs (human NG2+Olig2+, **Fig. 2G**), newly differentiated premyelinating oligodendrocytes (huNuclei+Bcas1+, **Fig. 2H)**, and mature oligodendrocytes (huNuclei+ASPA+) (**Fig. 2I**) exhibiting a typical myelinating morphology (**Fig 2J**). RNAscope confirmed the engraftment of human microglia-like cells (hCsf1r+) in the cerebral cortex, hippocampus, and meninges (**Fig. 2C**), and human astrocytes (hSlc1a3+) in the cerebral cortex (CTX), hippocampus (Hippo) and striatum (STR) (**Fig. 2B**). As shown in the quantification in **Fig. 2K-O**, consistently high and widespread engraftment of human glial cells (astrocytes, microglia-like cells and oligodendroglia) were observed in multiple brain regions, including the cortex, corpus callosum, and striatum. Although we did not directly transplant human microglia, Iba1+Csf1r+ human microglia-like cells were nonetheless detected in the xenografted mouse brains, likely originating from HSCs. The presence of microglia-like cells, coupled with the widespread engraftment of human astrocytes and oligodendroglia, establishes our model as a more comprehensive platform for neuroimmune research than traditional HSC transplantation-based humanized mouse models, which do not contain the full complement of human glia.

**Figure 2.**
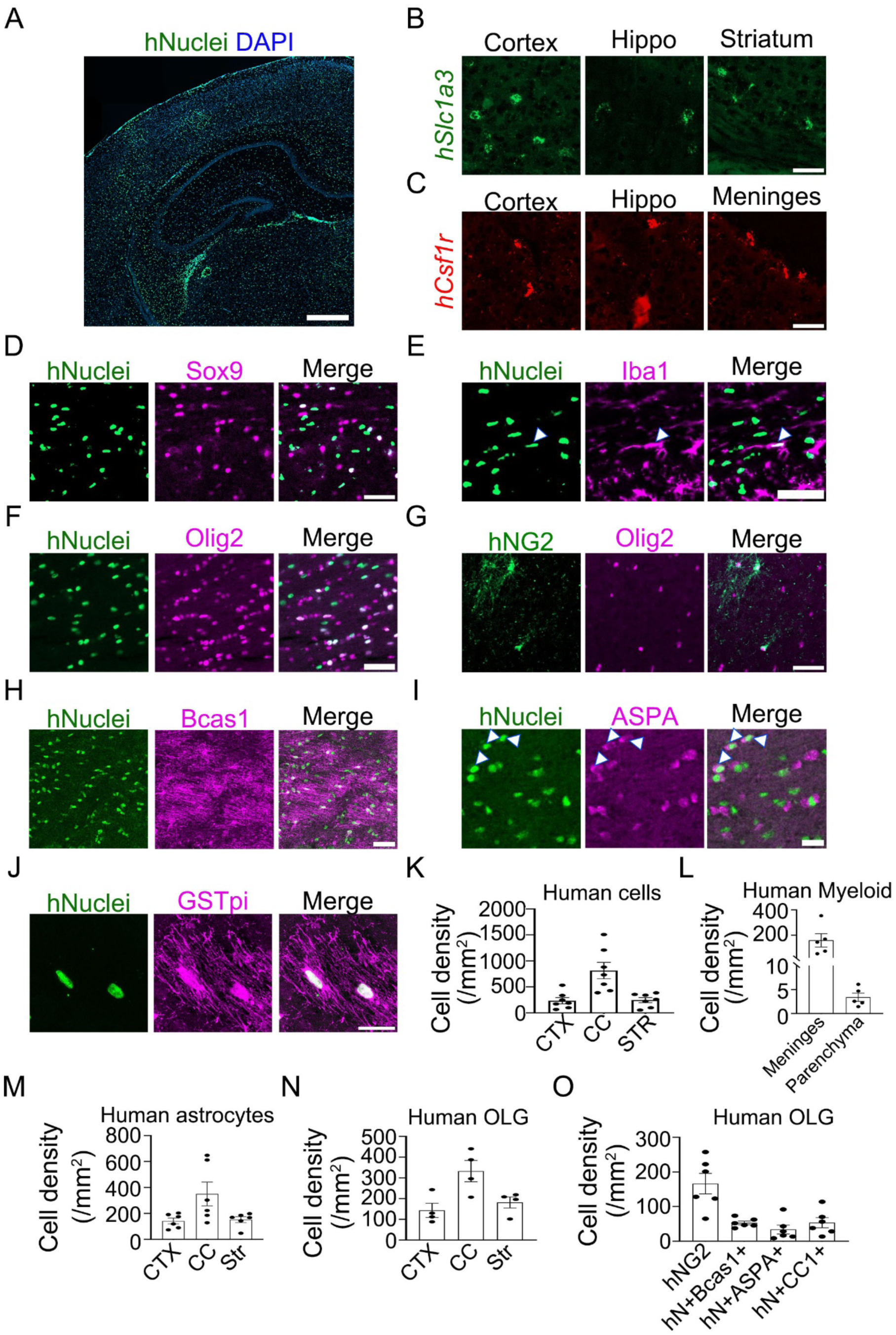
Widespread engraftment of human glia (astrocytes, OPCs, oligodendrocytes, and microglia-like cells) in the brains of glia-immune humanized mice. A) Human nuclei staining reveals widespread distribution of human cells in the chimeric mouse brain. Scale bar: 500 µm. Green: human nuclei. Blue: DAPI. B) *hSlc1a3* RNAscope assay demonstrates the presence of human astrocytes in the cortex, hippocampus, and striatum of the chimeric mouse brain. Scale bar: 50 µm. C) *hCsf1r* RNAscope assay shows human microglia-like cells in the cortex, hippocampus (hippo), and meninges of the chimeric mouse brain. Scale bar: 50 µm. D) Colocalization of human nuclei with astrocyte marker Sox9 indicates the presence of human astrocytes in the chimeric mouse brain. Scale bar: 50 µm. E) Colocalization of human nuclei with microglia-like cell marker Iba1 indicates the presence of human microglia-like cells (arrowheads) in the chimeric mouse brain. Scale bar: 50 µm. F) Colocalization of human nuclei with oligodendroglial marker Olig2 indicates the presence of Oligodendroglia in the chimeric mouse brain. Scale bar: 20 µm. G) Immunostaining of human NG2 shows the presence of human OPCs in the chimeric mouse brain. Scale bar: 50 µm. H) Colocalization of human nuclei with newly differentiated premyelinating oligodendrocyte marker Bcas1 indicates the presence of human immature oligodendrocytes in the chimeric mouse brain. Scale bar: 20 µm. I) Colocalization of human nuclei with mature oligodendrocyte marker ASPA indicates the presence of human mature oligodendrocytes in the chimeric mouse brain. Scale bar: 20 µm. J) Colocalization of human nuclear marker with the mature oligodendrocyte marker GSTpi confirms the presence of human oligodendrocytes with mature, myelinating morphology in the chimeric mouse brain. Scale bar: 20 µm. K) Quantification of the densities of human cells (human nuclei+ cells) in the cortex (CTX), corpus callosum (CC), and striatum (STR) (n= 7). L) Quantification of the densities of human meningeal myeloid cells (human *Csf1r*+ cells) and human microglia-like cells (human *Csf1r*+ cells) in the brain parenchyma of the chimeric mouse brain (n = 6). M) Quantification of the densities of human astrocytes (human nuclei+, Sox9+ double-positive cells) in the cortex, corpus callosum, hippocampus, and thalamus (Tha) of the chimeric mouse brain (n = 6). N) Quantification of the densities of human oligodendroglia cells (human nuclei+, Olig2+ double-positive cells) in the CTX, CC and Stratum (n = 4). O) Quantification of the densities of human OPCs (human NG2+ cells), human immature oligodendrocytes (human nuclei+, Bcas1+ double-positive cells), human mature oligodendrocytes (human nuclei+, ASPA+ double-positive cells), and human differentiated oligodendrocytes (human nuclei+, CC1+ double-positive cells) in the corpus callosum (n = 6).

### Robust HIV replication and immune activation in glia-immune humanized mice

To examine if glia-immune humanized mice support robust HIV infection and induce peripheral activation, mice were constructed and, following confirmation of successful immune reconstitution 14 weeks after transplantation, were either left uninfected or infected with the R5 tropic HIV-1_CH040_ via intravenous injection^17,18,25,26^. The levels of HIV replication over time were determined on plasma samples collected at multiple time points post-infection (Weeks 2, 4, 6, 8, and 10). The viral load in the plasma showed a progressive increase, with HIV RNA levels ranging from 1.2 × 10^6^ copies/ml at Week 2 to 6 × 10^6^ copies/ml at Week 10, demonstrating sustained viral replication over the 10-week observation period (**Fig. 3A**), similar to what was observed in the BLT humanized mouse model ^62^.

**Figure 3.**
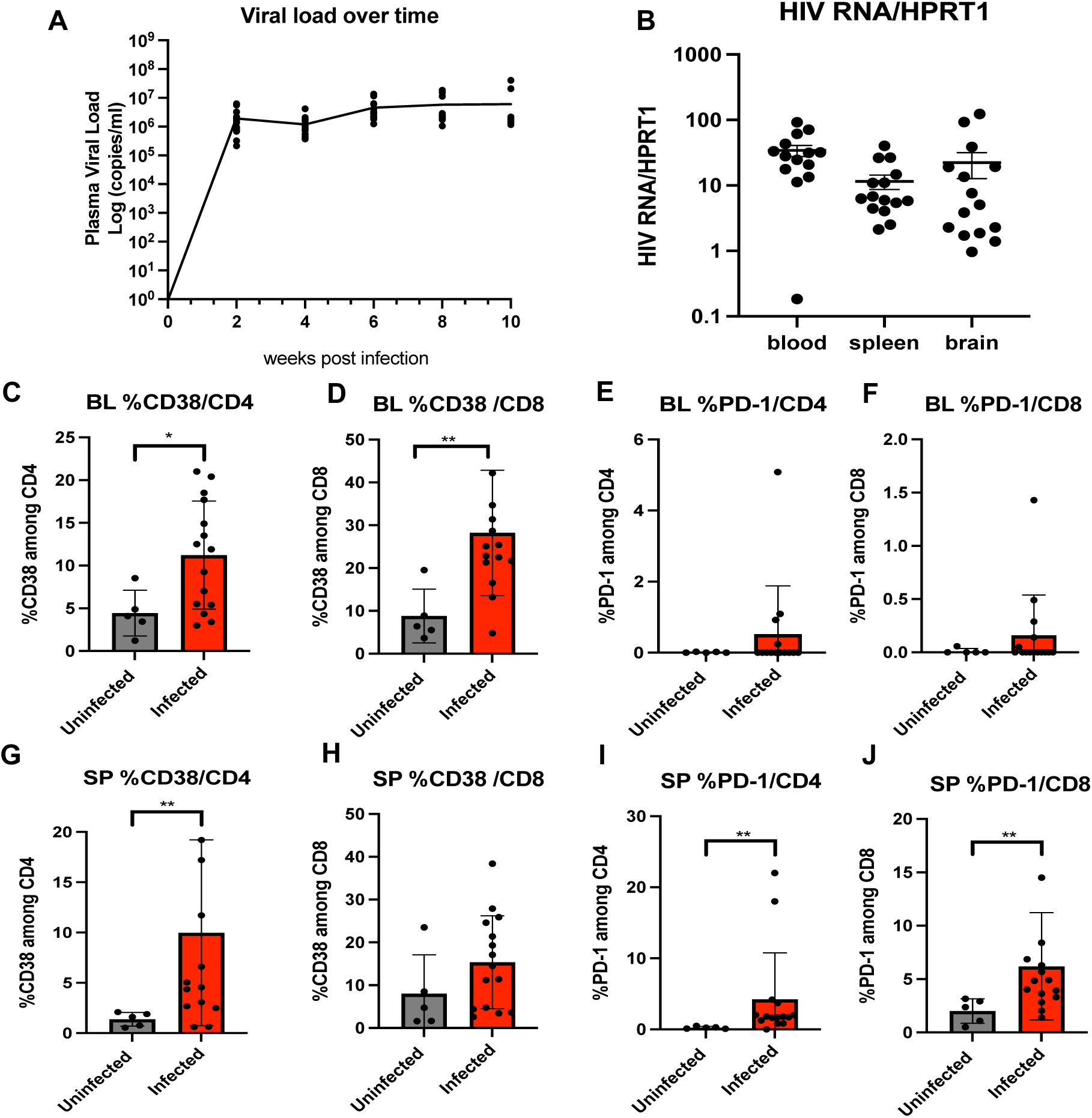
HIV infection and elevated Immune activation/exhaustion in infected glia-immune humanized mice. Glia-immune humanized mice were infected with HIV-1 CH040 (n=15) or mock infected (n=5) to evaluate viral replication and immune dysregulation. A) Plasma viral load was quantified longitudinally by real-time PCR. The X-axis represents time (in weeks) and the y-axis shows the plasma viral load in copies/ml (log scale). B) 10 weeks after HIV infection, mice were sacrificed, perfused with PBS and tissues were harvested. Viral RNA in blood, spleen and brain was quantified by real-time PCR. HPRT1 is internal control for normalization of gene expression. Data are presented as Mean± SEM. C-J) PBMCs from Blood (BL) and Splenocytes from spleen (SP) at week 10 were processed and stained for activation (CD38) (C, D, E, F) and exhaustion markers (PD-1) (G, H, I, J) on CD4^+^ and CD8^+^ T-cells using Flow-cytometry. Percentage of cells positive for each marker is represented as Mean ± S.D. * p=0.0328, **p=0.0037 in BL %CD38/CD8, **p=0.0077 in SP %CD38/CD4, **p=0.0024 in SP %PD-1/CD4, **p=0.0077 in SP %PD-1/CD8 by Mann-Whitney test.

In addition to plasma, we assessed viral replication in lymphoid and CNS tissues. 10 weeks after HIV infection, mice were euthanized and perfused with phosphate buffered saline (PBS), with the left hemisphere of the brain fixed for IHC and RNAscope, while the right hemisphere was processed for flow cytometry, real time PCR, and RNAseq analysis. HIV RNA was detected in the blood, spleen, and PBS-perfused brain tissues using quantitative real-time PCR. The data revealed substantial HIV RNA expression in all three tissues, with cycle threshold (Cq) values ranging from 26.34 to 42.20 in the blood, 20.18 to 30.12 in the spleen, and 26.46 to 33.37 in the brain. As shown in **Fig 3B**, the HIV-RNA analysis was normalized to the expression level of housekeeping gene HPRT1. These findings confirm robust and widespread HIV replication in both lymphoid tissues and the brain, indicating the model’s suitability for studying HIV dynamics in peripheral and CNS compartments.

HIV infection-mediated chronic inflammation is well documented in PLWH and in infected humanized mice ^24,27,63^. To assess whether glia-immune humanized mice exhibit HIV infection-induced chronic inflammation, we used flow cytometry to analyze the expression of key activation marker CD38 and exhaustion marker PD-1 on T cells from uninfected and infected glia-immune humanized mice. Activation marker CD38 and exhaustion marker PD-1, were both markedly increased on CD4+ and CD8+ T cells in the blood (**Fig 3C-F**) and the spleen (**Fig 3G-J**) of HIV infected mice as compared to uninfected mice. These findings demonstrate that glia-immune humanized mice recapitulate key features of HIV pathogenesis, including robust viral replication in peripheral and CNS compartments and chronic peripheral immune activation, supporting the model’s relevance for studying systemic and neuroimmune consequences of HIV.

### HIV infection elevates type I IFN responses, and the “immune oligodendroglia” state in the CNS

HIV infection induces type I IFN responses in PLWH ^64,65^. We and others have shown that chronic type I IFN activation leads to chronic inflammation, T cell exhaustion, and maintenance of HIV reservoirs ^9,25,48^. To examine whether glia-immune humanized mice exhibit type I INF response, a key mechanism for HIV pathogenesis, we examined type I IFN response gene (ISG) MX1 expression by real-time PCR. We observed that infected mice showed significantly elevated levels of MX1 in blood, spleen and brain tissues compared to uninfected controls (**Fig. 4A**), mirroring observations in PLWH^66–68^ and reflecting the ongoing immune activation in response to persistent HIV replication. To identify molecular changes in human glia in response to HIV infection, we performed bulk RNA-seq and analyzed transcripts mapped to the *human* reference genomes using brain samples from HIV-infected versus uninfected mice. RNA-seq revealed a striking induction of type I ISGs and associated antiviral pathways in human glial cells from infected mice (**Fig. 4B-D**). Differential gene expression analysis (Volcano plot in **Fig. 4B**) identified multiple significantly upregulated ISGs (e.g., OAS3, ISG15, IFI44, IFI44L, IFIT3) alongside classical interferon-regulatory factors (IKZF2, IFNGR1), suggesting a heightened antiviral state in the HIV-infected brain. Network analysis of differentially expressed genes (**Fig. 4C**) highlighted a large, interconnected module of upregulated genes (red) enriched for interferon-associated factors, whereas a cluster of downregulated genes (blue) encompassed components implicated in transcriptional regulation (ZNF652, IKZF2), metabolism (ACSS2, DUT, STIM1) and epigenetic control (EP300). Consistent with these observations, gene set enrichment analysis (**Fig. 4D**) demonstrated that upregulated genes were significantly overrepresented in pathways related to “Interferon alpha/beta signaling”, “PKR-mediated signaling”, and “ISG15 antiviral mechanisms”. Immunohistochemistry (IHC) further confirmed increased IFN-I response gene human MX1 expression in the brains of infected mice (**Fig. 4E-I**). We observed increased density of hMX1+ glial cells in corpus collosum, cortex, hippocampus and thalamus of the glia immune humanized mice (**Fig.4G-I**). These results reveal molecular changes induced by HIV infection in human glia in vivo, further demonstrating the value of the glia-immune humanized mouse model for investigating the molecular mechanisms of HIV CNS infection and other neurological disorders.

**Figure 4:**
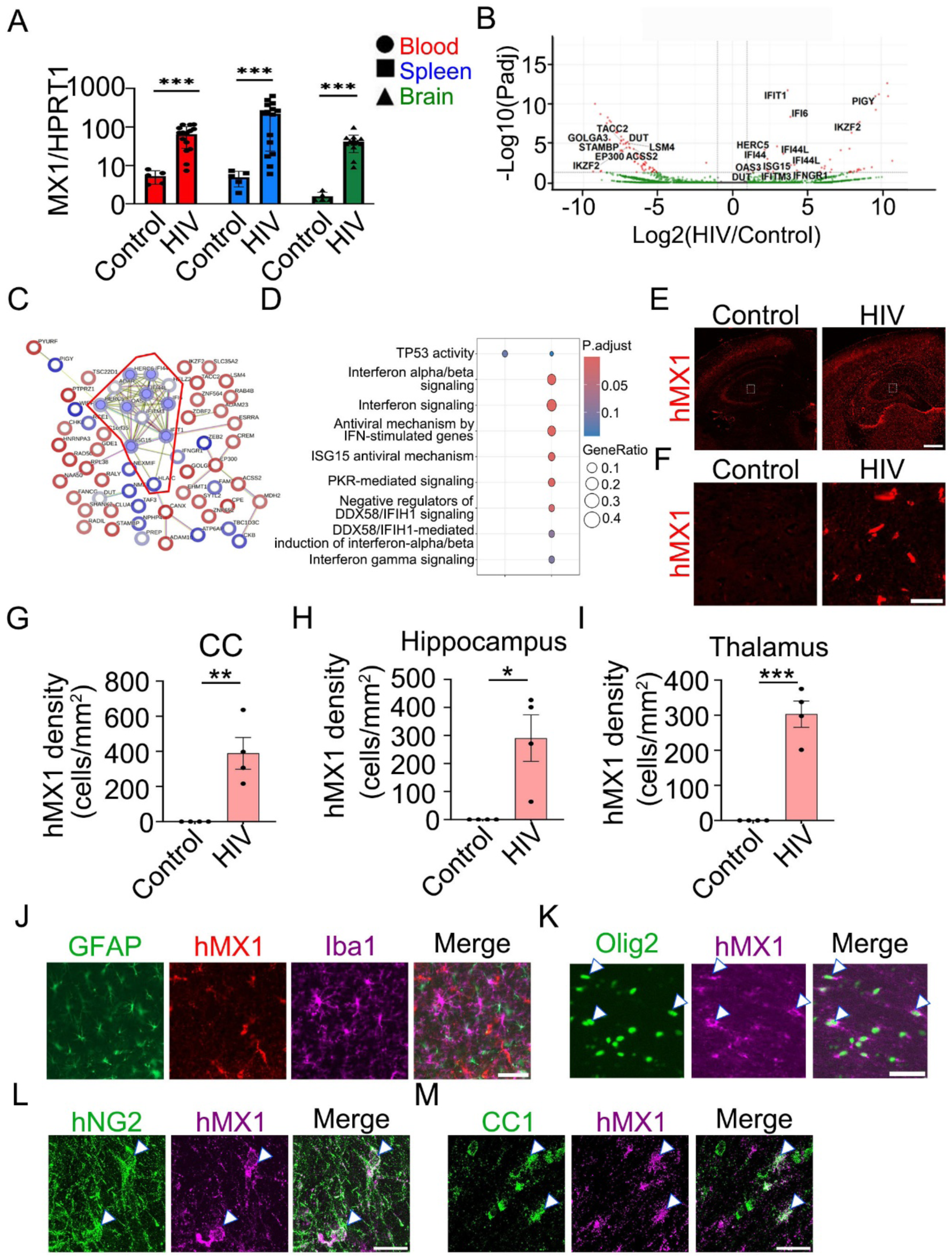
HIV infection induces upregulation of human-specific interferon response gene MX1 in glia-immune humanized mice. A) Interferon stimulated gene-MX1 expression level in blood, spleen and brain as measured by real-time PCR in uninfected (n=5) and infected mice (n=15). HPRT1 is internal control for normalization of gene expression. B) Volcano plot showing differential gene expression of infected versus uninfected brains. The x-axis represents the log₂ fold change (log₂FC) in gene expression while the y-axis shows the −log₁₀ p-value. Genes with significant differential expression are highlighted in red. C) Reactome pathway analysis of significantly differentially expressed genes in the brains of infected versus uninfected mice. This network visualization displays the interactions between genes with significant changes in expression. Nodes represent individual genes, and edges indicate known or predicted interactions based on pathway data. The network shows a highly interconnected cluster of genes, including ISG15 and OAS1 involved in interferon signaling pathways and the antiviral response. D) Gene enrichment analysis. Red circle highlights genes involved in interferon signaling. E) Human MX1 (hMX1) staining shows a widespread upregulation of the interferon response in the brains of glia-immune humanized mice following HIV infection (n=4). Scale bar: 500 µm. Zoomed in images (dashed rectangular area in F) shows increased human MX1+ cells in the hippocampus of the glia-immune humanized mouse. (G-I). Quantification of the densities of human MX1+ cells in the corpus callosum (CC), cortex, hippocampus, and thalamus of the chimeric mouse brain. (*p=0.0127 by unpaired t-test in Hippocampus, **p=0.0051 by unpaired t-test in CC, ***p=0.0002 by unpaired t-test in Thalamus, ***p=0.0003 in blood, ***p=0.0005 in spleen, **p=0.0001 in brain by Mann-Whitney test.) J) Limited colocalization of hMX1 (red) with astrocyte (GFAP+ green) or microglia (Iba1+ purple). K) Colocalization (yellow) of hMX1 (red) with Oligodendroglia (Olig2+ in green). L) Colocalization (white) of hMX1 (purple) with human oligodendroglia precursor cells (hNG2+ in green). Scale bar: 20 µm. M) Colocalization (white) of hMX1 (purple) with differentiated oligodendroglia marker (CC1+ in green). Scale bar: 20 µm.

To identify which CNS cell type(s) upregulate MX1 expression during HIV infection, we performed co-immunostaining of MX1 with Iba1 (microglia-like cells), GFAP (astrocytes), Olig2 (OPCs and oligodendrocytes), NG2 (OPCs), and CC1 (oligodendrocytes) (**Fig.4 J-M**). Surprisingly, we observed more extensive colocalization of MX1 with Olig2, NG2, and CC1 (**Fig.4K-M**), whereas colocalization with Iba1 or GFAP was minimal (**Fig 4J**). These findings suggest that oligodendroglia are the predominant cell type upregulating MX1 expression in the HIV-infected brain. To assess potential regional heterogeneity in MX1 expression among oligodendroglia, we quantified the percentage of MX1+ cells among human oligodendroglia across different brain regions. We found a lower proportion of MX1+ oligodendroglia in the corpus callosum compared to the cortex and striatum (**Supplementary Fig. 1**), suggesting that oligodendroglia from distinct brain regions differ in their susceptibility to adopting the immune oligodendroglia state during HIV infection.

Interestingly, we observed a significant increase in ASPA+Olig2+ mature oligodendrocytes in the corpus callosum (**Fig 5 A, C, D, F**) and striatum (**Fig 5G**). However, this increase was only evident when both human and mouse cells were included in the quantification, whereas the densities of huNG2+ (human OPCs), huNuclei+ASPA+ (human mature oligodendrocytes) or huNuclei+Olig2+ cells (all human oligodendroglia) were unchanged (**Supplementary Fig. 2**). In contrast to the increase in mature oligodendrocytes, densities of Bcas1+ immature oligodendrocytes remained unchanged (**Fig. 5B, E**).

**Figure 5.**
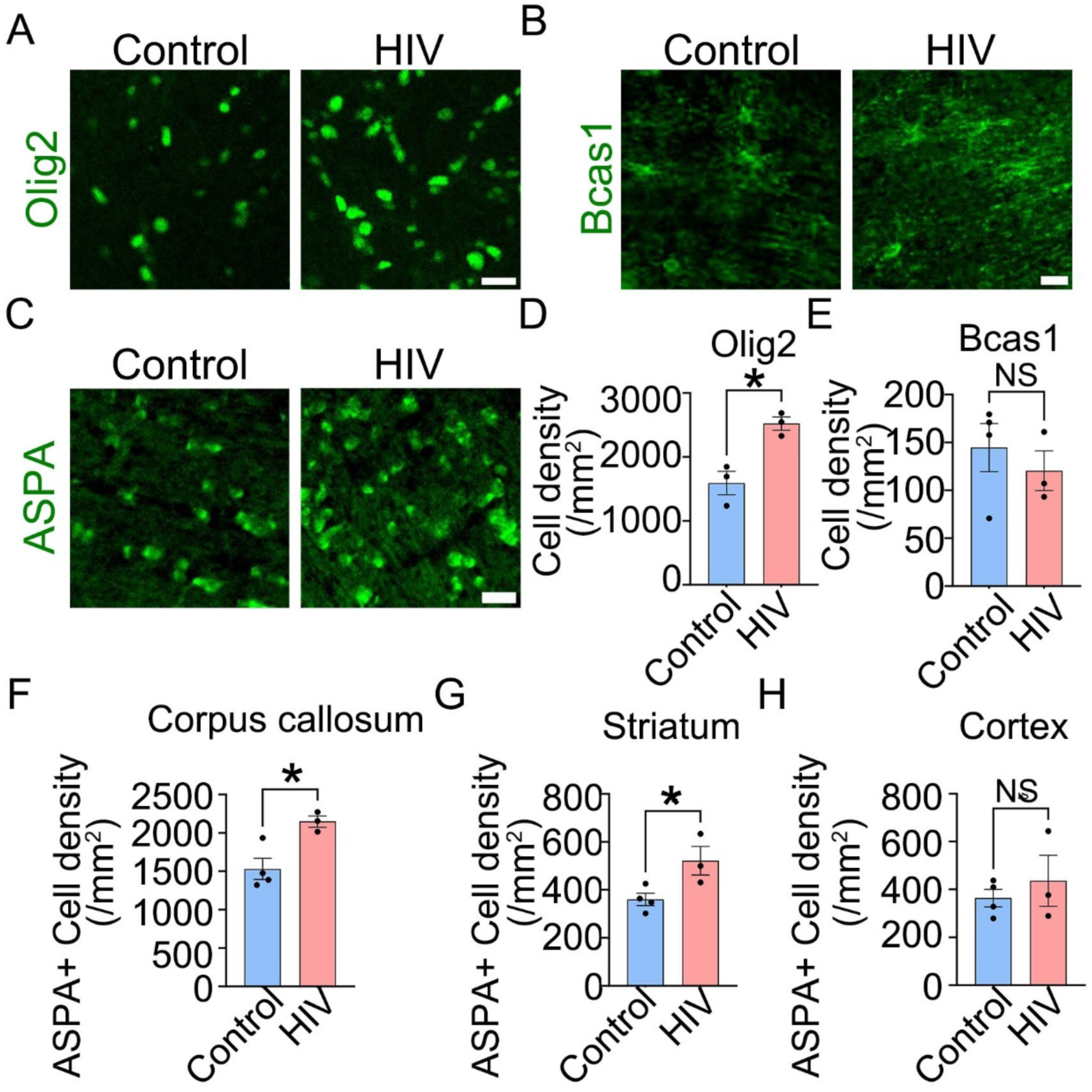
HIV infection increases the densities of oligodendrocytes in glia-immune humanized mice. A) Immunostaining of Olig2 shows an increase in the number of oligodendroglia (olig2+ cells) after HIV infection. Scale bar: 20 µm. B) Immunostaining of Bcas1 shows no change in the number of immature oligodendrocytes (Bcas1+ cells) after HIV infection. Scale bar: 20 µm. C) Immunostaining of ASPA shows an increase in the number of mature oligodendrocytes (ASPA+ cells) after HIV infection. Scale bar: 20 µm. D) Quantification of the densities of oligodendroglia (Olig2+ cells) in the corpus callosum of the chimeric mouse brain. (n = 3, p = 0.0117 by unpaired t-test.) E) Quantification of the densities of immature oligodendrocytes (Bcas1+ cells) in the corpus callosum of the chimeric mouse brain. (n = 4 for Control, n = 3 for HIV, p = 0.5128, unpaired t-test). F to H) Quantification of the densities of mature oligodendrocytes (ASPA+ cells) in the corpus callosum, striatum, and cortex of the chimeric mouse brain. (n = 4 for Control, n = 3 for HIV, p = 0.0169 for CC, p = 0.0395 for STR, p = 0.4962 for CTX, unpaired t-test).

Consistent with our observations, studies of postmortem patient tissue show elevated Interferon-mediated signaling pathway in OPCs from HIV infected individuals^39,69^. Oligodendrocyte density was shown to be elevated in patients with mild pathology while decreased in untreated patients who died of AIDS^70,71^. Although oligodendroglia has traditionally been studied exclusively for their role in forming myelin, emerging evidence indicates that OPCs and oligodendrocytes can acquire an immunomodulatory phenotype. Here, our observation supports the presence of “immune OPCs” and “immune oligodendrocytes” that have elevated IFN-I gene expression in HIV infected brains, highlighting underappreciate roles of oligodendroglia in HIV CNS pathology.

### HIV infection leads to microglia and astrocyte reactivity and loss of synaptic proteins in glia-immune humanized mouse

To determine if HIV infection disturbs brain homeostasis, we examined the status of microglia and astrocytes in these mice. As key regulators of central nervous system homeostasis, microglia and astrocytes function as the brain’s primary immune responders. Increases in microglia marker Iba1 and reactive astrocyte marker GFAP are often associated with neuroinflammation and neuropathology ^72^ ^73^. We quantified microglia density (Iba1+ cells) (**Fig. 6A**) and reactive astrocyte density (GFAP+ cells) (**Fig. 6B**) in four brain regions: the cortex, corpus callosum, hippocampus, and thalamus. Microglia density significantly increased in the hippocampus and thalamus compared to control (**Fig. 6C-F**). Similarly, the density of reactive astrocytes also increased in the corpus callosum and hippocampus compared to control (**Fig. 6G-J**).

**Figure 6.**
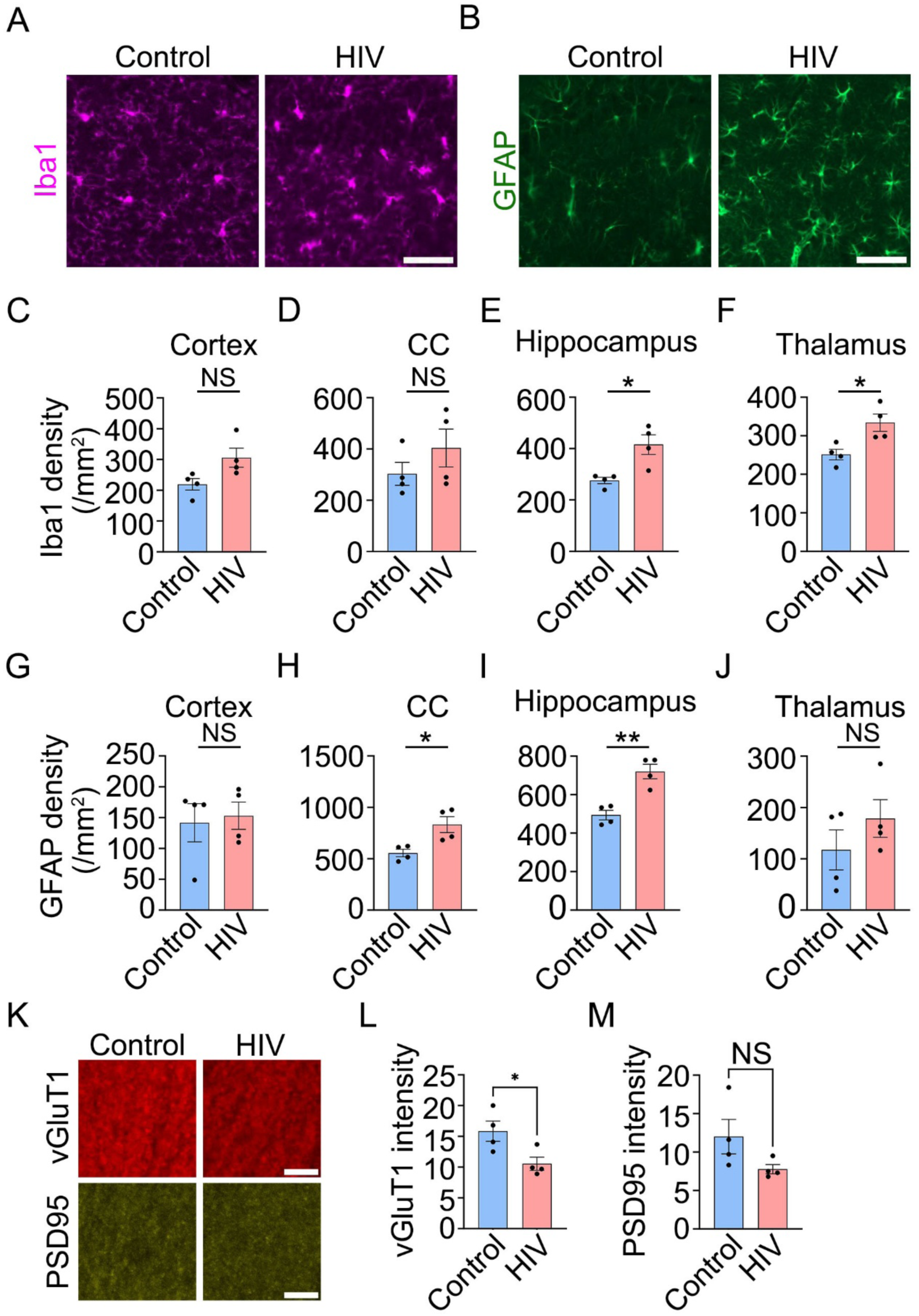
HIV infection leads to microglia, astrocyte reactivity and loss of synaptic proteins in the chimeric mouse brain. A) Iba1 staining in the hippocampus shows that microglia density is increased after HIV infection. Scale bar: 50 µm. B) GFAP staining in the hippocampus shows that GFAP+ astrocytes density is increased after HIV infection. Scale bar: 50 µm. (C - F) Quantification of the density of Iba1+ cells in the cortex, corpus callosum, hippocampus, and thalamus of the chimeric mouse brain. (n = 4, p = 0.0553 for CTX, p = 0.288 for CC, p = 0.0131 for Hippo, p = 0.0197 for thalamus). (G - J) Quantification of the density of GFAP+ cells in the cortex, corpus callosum, hippocampus, and thalamus of the chimeric mouse brain. (n = 4, p = 0.7771 for CTX, p = 0.0182 for CC, p = 0.0024 for Hippo, p = 0.2966 for thalamus). (*p<0.05, **p<0.005, by unpaired t-test). (*p<0.05, **p<0.005, by unpaired t-test). K) vGluT1 and PSD95 staining in the CA1 region of hippocampus shows a decrease in vGluT1 immunofluorescence intensity after HIV infection and no change in PSD95 immunofluorescence intensity. Scale bar: 5 µm. L) Quantification of the intensity of vGluT1 in the CA1 region of hippocampus from the chimeric mouse brain. (n = 4, p = 0.0355). M) Quantification of the intensity of PSD95 in the CA1 region of hippocampus from the chimeric mouse brain. (n = 4, p = 0.1178 for CTX).

Synapse loss is a hallmark of HAND, and its extent correlates with cognitive impairment. To evaluate synaptic changes, we conducted IHC for the presynaptic protein VGLUT2 and the postsynaptic protein PSD-95. In the CA1 region of the hippocampus, HIV-infected mice exhibited a significant reduction in VGLUT2 immunofluorescence compared to controls (**Fig. 6K&L**), whereas PSD-95 immunofluorescence remained unchanged (**Fig. 6K&M**). These results suggest that our humanized mouse model recapitulates HIV infection-associated synapse loss in patients, providing a valuable model for mechanistic study and drug testing for HIV associated cognitive impairment.

## Discussion

Previous humanized mouse models, which typically incorporate a single type or a limited subset of human cells, have been instrumental for preclinical studies of disease mechanisms^7,48,74–87^. Building on these foundations, the next-generation glia-immune humanized mouse model described here represents an important advancement, enabling in vivo investigation of human neuroimmune interactions with greater cellular complexity and physiological relevance. By integrating all four major types of human glial cells-astrocytes, microglia-like cells, OPCs, and oligodendrocytes - alongside donor-matched peripheral immune cells within the same mouse, this model recapitulates the complex cellular landscape and intercellular interactions of the human CNS more faithfully than previous systems. For example, glial cells present antigen but HLA-mismatched antigen presentation cells cannot properly engage with T cells and may themselves be subject to immune rejection. Thus, establishing a humanized mouse model incorporating HLA-matched, full complement of human glial and immune cell types would provide an indispensable platform for human neuroimmune interaction studies.

Here, we demonstrated the value of this model for CNS HIV research. Current humanized mouse models, while invaluable for studying peripheral HIV infection, have been limited in their utility to understand CNS infection due to the lack of matched human glia in the brain. Glia, including microglia, astrocytes, OPCs, and oligodendrocytes, are essential mediators of CNS homeostasis and immune responses, and their dysfunction in the setting of HIV infection is implicated in neurological complications ^13,88,89^. By incorporating donor-matched glial cells, our model supports robust engraftment of astrocytes, OPCs, oligodendrocytes, and microglia-like cells, and simultaneous robust peripheral immune reconstitution of donor matched HSCs. Our model provides a unique platform to investigate HIV infection that spans peripheral organs and the CNS.

Our data demonstrated that our model supports robust and persistent HIV replication in the peripheral blood, lymphoid tissues and in the brain. The elevated proinflammatory gene expression in both the periphery and brain underscores the potent immunopathological effects of HIV infection. Specifically, RNA sequencing of brain tissues demonstrated upregulation of type I ISGs and inflammasome-associated genes in human glial cells. Importantly, we observed glia reactivity in HIV infected animals, including in oligodendroglia and astrocytes, which are not key reservoirs of HIV, suggesting that bystander reactivity of glial cells may play key role in CNS inflammation and HAND^90^. Interestingly, in the brains of infected glia-immune humanized mice, we also observed several down regulated genes that are related to glia metabolism. For example, STIM1 is involved in calcium signaling, which is essential for astrocyte and microglial responses to inflammatory stimuli^91,92^. Downregulation of STIM1 suggests an impairment in calcium signaling pathways, potentially leading to glial dysfunction and contributing to neuroinflammation^93^. Similarly, decreased ACSS2 levels could affect neuronal energy metabolism and histone acetylation, impacting neuronal function and memory formation^94–96^. Therefore, our model offers a powerful platform to investigate the molecular mechanisms of HIV-driven neuroinflammation and glial dysfunction in the brain.

Our discovery that human OPCs and oligodendrocytes adopt an immune phenotype - marked by robust upregulation of type I ISG MX1 expression in HIV-infected glia-immune humanized mouse brains provides novel insights into the dynamic, immunomodulatory potential of human oligodendroglia in the diseased CNS. These results complement reports of oligodendroglia damage in HIV infection^13,14,97–99^. Traditionally viewed primarily as myelinating cells and precursors, these findings position oligodendroglia as active participants in neuroimmune responses, capable of sensing inflammatory cues and engaging in immune signaling. Although immune oligodendroglia have been described in models of multiple sclerosis and aging^100–102^, their presence in the context of HIV infection has not, to our knowledge, been previously reported. In HIV infection, this immune oligodendroglia state may contribute to neuroinflammation and white matter damage in synergy with canonical immune cells such as microglia and infiltrating macrophages and T cells. These observations raise the possibility that oligodendroglia may serve as amplifiers or modulators of CNS inflammation in HAND and potentially in other neuroinflammatory and/or demyelinating disorders. Future research should explore whether this immune oligodendroglia phenotype is reversible, whether/how it contributes to impaired remyelination or neural circuit dysfunction, and how it is regulated by specific inflammatory mediators or viral products. Our model provides a novel platform for investigating the function of human immune oligodendroglia in vivo in a range of neurological disorders and our findings point to human oligodendroglia as a previously untapped therapeutic target in neuroimmune disorders.

The ability of this glia-immune humanized mouse model to recapitulate HIV infection and inflammation in the CNS offers a valuable tool for testing curative and neuroprotective interventions. Our model supports a full range of biochemical, imaging, and physiological techniques, creating a robust platform for translational research on neuro-immune interactions in HIV CNS comorbidities and other CNS diseases. In addition, throughout the seven-month study period, we did not observe graft-versus-host disease (GVHD) in our glia-immune humanized mice, highlighting the stability and tolerability of the dual transplant methods.

The survival of myeloid cells requires species-specific M-CSF^103^. Thus, previous methods for reconstituting human myeloid cells in mice typically require transgenic expression of human M-CSF^75,80–82^. In our study, neonatal transplantation of human HSCs and glia is sufficient to maintain the survival of some human myeloid cell in the blood and brain, suggesting that either the transplanted human cells and their progeny produce M-CSF or the neonatal host mouse provides alternative growth factors that compensate for the species-specific need for human M-CSF. In future studies, utilizing host mice that express human M-CSF to construct glia-immune humanized models may further enhance the engraftment and expansion of human myeloid cell populations.

In summary, our glia-immune humanized mouse model fills a critical gap in HIV research by integrating donor-matched human glia and a functional human immune system. Our findings highlight the capacity of these mice to support HIV replication within both peripheral and CNS tissues, mirroring key features of human HIV neuropathogenesis. Beyond HIV research, such humanized mouse models hold tremendous potential for advancing our understanding of other neurological diseases that involve complex interactions between human glia and immune cells. There are many major differences between human and mouse immunology in both innate and adaptive immune responses^104^, and in glial cells^3,79^. Our model will enable the mechanistic dissection of neuroimmune signaling pathways, human glial-immune crosstalk, and their roles in disease initiation and progression across a spectrum of neurological disorders, including neuroinfectious diseases, demyelinating diseases, neurodegenerative disorders, and neuroinflammatory conditions. Furthermore, this model provides a unique opportunity for the development and pre-clinical testing of human immune therapies (such as antibody-based therapeutics ^105^ or Chimeric Antigen Receptor therapy^106^) that modulate neuroimmune interactions in CNS disorders. In future work, deeper single-cell analyses and spatial transcriptomic approaches can further delineate the molecular interaction among astrocytes, OPCs, oligodendrocytes, microglia, neurons, and peripheral immune cells during HIV infection.

## Materials and Methods

### Isolation of human astrocytes and OPCs

All mid-gestation tissue-related experiments were carried out in accordance with relevant federal, state, and local laws, including adherence to required regulatory approvals and informed consent procedures. Human glia were isolated from gw 18 brains using a protocol adapted from a rat glia enrichment protocol^107^. Briefly, gw 18 brains were chopped into small pieces using a razor blade and enzymatically dissociated with papain. Dissociated cells were plated in a T75 flask and maintained in a serum-containing culture medium. Three days later, unwanted cells such as neurons were depleted by mechanical shaking. Remaining attached cells, which contains both astrocytes and OPCs, were maintained in culture until transplantation.

### Generation of glia-immune humanized mouse

We developed a novel glia-immune humanized mouse model, where the mice were reconstituted with donor-matched human glia in the brain and a human immune system in the peripheral and lymphoid tissues. P2-6 immunodeficient NSG mice (NOD.Cg-Prkdc^scid Il2rg^tm1Wjl/SzJ) were injected with 0.5 million donor-matched, gestation <18 weeks liver-derived CD34+ HPSCs via intrahepatic injection and 400,000 brain-derived astrocytes and oligodendroglial cells via intracranial injection at four sites in the subcortical regions of the brain, including the striatum and thalamus bilaterally^3^. Fourteen weeks post-reconstitution, successful immune reconstitution was confirmed in the peripheral blood of the humanized mice, with detectable levels of human T cells, B cells, and monocytes.

### Infection of mice and quantification of viral loads

Glia-immune humanized mice were inoculated with HIV-1 CH040 ^108^ (500 ng of p24 per mouse) via retro-orbital injection under general anesthesia. To assess HIV plasma viremia every two weeks, viral RNA was extracted from plasma samples, and a one-step real-time PCR was conducted using the TaqMan RNA-to-Ct 1-Step kit (Thermo Fisher Scientific, USA). PCR amplification was performed with the following primers and probe:

CH040 forward primer: CAATGGCAGCAATTTTACCA

CH040 reverse primer: GAATGCCGAATTCCTGCTTGA

CH040 dye-labeled probe: 5’ [FAM]-CCCACCAACAGGCGGCCTTCACTG-[NFQ]-3’

### Nucleic acid isolation and real time PCR

To assess the expression levels of the Type 1 interferon (IFN) response gene, MX1, and cell-associated HIV RNA, RNA was extracted from harvested cells following the manufacturer’s protocol (Qiagen). cDNA was synthesized using the High-Capacity cDNA Reverse Transcription Kit (Thermo Fisher Scientific, USA). For the quantification of MX1 and HPRT1, TaqMan Gene Expression Assays (Thermo Fisher Scientific, USA) were used with the following probes: Human MX1 (Hs00895608_m1) and Human HPRT1 (Hs01003267_m1). For HIV RNA quantification, the primers and probes specific to the CH040 strain, as described above, were utilized.

### Flow cytometry

Single-cell suspensions prepared from the peripheral blood and spleen of glia-immune humanized mice were stained with the following antibodies: CD45 (clone HI30), CD3 (clone OKT3), CD4 (RPA-T4), CD8 (RPA-T8), CD14 (clone 61D3), CD11c (clone 3.9), CD11b (clone ICRF44), CD19 (HIB19), PD-1 (clone EH12.2H7), and CD38 (clone HIT2). To facilitate live cell gating, the LIVE/DEAD Fixable Yellow Dead Cell Stain Kit (Invitrogen, USA) was used. Antibodies for cell surface markers were conjugated to a range of fluorophores, including BV785, BV605, Pacific Blue, Alexa Fluor 700 (AF700), Phycoerythrin (PE), ECD, PE-Cy7, Allophycocyanin (APC), PerCP-Cy5.5, and PE-Cy5 in appropriate combinations. Data acquisition was performed using the LSR Fortessa flow cytometer and FACS Diva software (BD Biosciences), while FlowJo software (version 10.7.2) was used for data analysis.

### IHC

Glia-immune humanized mice were anesthetized with isoflurane and transcardially perfused with 1× PBS (pH 7.4), followed by fixation with 4% paraformaldehyde (PFA). Brains were post-fixed in 4% PFA at 4 °C overnight, then transferred to a 30% sucrose solution at 4 °C for two days or until they sank. Samples were embedded in optimal cutting temperature (OCT) compound (Fisher, cat#23-730-571) and stored at −80 °C until sectioning. Brain tissue was cut into 30 μm slices using a Leica cryostat and stored in cryoprotectant solution (30% sucrose, 1% polyvinylpyrrolidone, and 30% ethylene glycol in 0.1 M phosphate buffer) at −20 °C. For immunostaining, sections were blocked and permeabilized with 10% donkey serum and 0.3% Triton X-100 in PBS, then incubated with primary antibody solutions overnight at 4 °C. For human NG2 and Bcas1 staining, primary antibody incubation was extended to 2 days. Heat-induced antigen retrieval was performed to reveal ASPA signals. Sections were first rinsed with antigen retrieval buffer (10 mM sodium citrate, 0.05% Tween-20, pH 6.0) for 5 min at room temperature. They were then transferred into pre-heated antigen retrieval buffer and subjected to heat-induced antigen retrieval using a vegetable steamer for 10 min. After heating, sections were allowed to cool to room temperature in the same buffer before proceeding to blocking and immunostaining. The following primary antibodies were used: anti-human nucleus protein (Chemicon, cat#MAB1281, 1:500 dilution), anti-human nuclei (Thermo Fisher, cat# PA5-18498, 1:500 dilution), anti-Iba1 (Wako, cat#019-19741, 1:500 dilution), anti-Olig2 (Millipore, cat#211F1.1, 1:1000 dilution), anti-Sox9 (R&D, cat#AF3075, 1:1000 dilution), anti-GFAP (Dako, cat# Z0334, 1:1000 dilution), anti-human MX1 (Abcam, cat#ab284603, 1:500 dilution), anti-Bcas1 (Synaptic system, cat# 445 003, 1:250 dilution), anti-ASPA (Genetex, cat# GTX113389, 1:500 dilution), Anti-GST-π (MBL International corporation, cat#312, dilution 1:500), anti-human NG2 (Sigma, cat# MAB2029, 1:200 dilution), anti-vGluT1 (Synaptic system, cat#N1602-AF647-L, 1:500 dilution), and anti-PSD95 (Thermo Fisher, cat#51-6900, 1:500 dilution). After primary antibody incubation, sections were washed three times with PBS and incubated with secondary antibodies at room temperature for 2 hours, followed by three additional washes in PBS. Stained sections were mounted onto Superfrost Plus slides (Fisher, cat#12-550-15), covered with mounting medium (Fisher, cat#H1400NB), and sealed with glass coverslips. Fluorescent imaging was carried out on a Zeiss Apotome epifluorescence microscope and a Leica Stellaris confocal microscope with consistent power and exposure settings for all samples processed with the same probe set.

### RNAscope

Glia-immune humanized mice were transcardially perfused with RNase-free PBS followed by fixation with RNase-free 4% PFA. Brains were removed and post-fixed in 4% PFA for an additional 2 hr at room temperature and then overnight at 4 °C. Brains were dehydrated in RNase-free 30% sucrose, embedded in OCT compound, and cut into 20 μm-thick sections using a Leica cryostat. RNAscope experiments were performed using the RNAscope Multiplex Fluorescent Reagent Kit v2 (ACDBio, cat#323100) according to the manufacturer’s protocol. Probes specific to human SLC1A3 (cat#461081-C1), and human CSF1R (cat#310811-C2) were obtained from ACDBio and used for this study. Fluorescent imaging was carried out on a Zeiss Apotome epifluorescence microscope.

### RNAseq

Total RNA was extracted from humanized mouse brain tissues and prepared for Illumina sequencing following standard protocols. Technical replicate FASTQ files were concatenated, and initial quality assessments were performed using FastQC. Unique molecular identifiers (UMIs) were extracted with UMI-tools, while adapter and low-quality bases were trimmed with Trim Galore!. We then removed genomic contaminants using BBSplit and filtered out rRNA sequences with SortMeRNA. To accommodate reads derived from both mouse and human cells, we constructed a mixed reference by combining Gencode GRCm39 (mouse) and GRCh38 (human) assemblies and annotations, creating a single index for genome alignment. Reads were aligned to this hybrid reference using STAR, followed by transcript quantification in alignment-based mode with Salmon. Aligned reads were sorted and indexed (SAMtools), and UMIs were used for deduplication with UMI-tools, complemented by duplicate marking (picard MarkDuplicates). We then assembled transcripts using StringTie, and coverage tracks (bigWig format) were generated via BEDTools and bedGraphToBigWig. Multiple quality-control checks were performed (RSeQC, Qualimap, dupRadar, and Preseq) to evaluate alignment, duplication rates, and library complexity. Finally, differential expression was assessed using DESeq2, and downstream functional analyses—including Gene Ontology, Reactome pathway enrichment, and additional annotation through clusterProfiler—were conducted in R.

### Study approval

Peripheral blood mononuclear cells were obtained at UCLA in accordance with UCLA IRB–approved protocols under written informed consent using an IRB-approved written consent form by the UCLA Center for AIDS Research Virology Laboratory and distributed for this study without personal identifying information. Human tissue was obtained from UCLA Pathology Core Laboratory without identifying information and did not require IRB approval for its use. Animal research described in this article was performed under the written approval of the UCLA Animal Research Committee in accordance with all federal, state, and local guidelines. All surgeries were performed under ketamine/xylazine and isoflurane anesthesia, and all efforts were made to minimize animal pain and discomfort.

## Acknowledgement

We thank UCLA Center for AIDS Research (CFAR) Humanized Mouse Core staff research associate Nianxin Zhong for his assistance in the humanized mice work. This work was funded by the National Institute of Allergy and Infectious Diseases (R01AI172727 to AZ and MDM), the National Institute on Drug Abuse (R01DA052841, R01DA059873 to AZ), National Institute of Health grants P30AI28697 (to the UCLA CFAR Virology Core, Gene and Cell Therapy Core, and Humanized Mouse Core) and U19AI149504 (to SGK and Dr. Irvin Chen [UCLA]); the California Institute for Regenerative Medicine (grant TRAN1-14625 to SGK.); California HIV/AIDS Research Program (grant H24BD7817 to WM); UCLA-Charles Drew University (CDU) CFAR (grant AI152501 to WM); the NIH/NINDS R01NS109025, the Broad Stem Cell Research Center (BSCRC) Innovation Award, the W.M. Keck Foundation Junior Faculty Award, UCLA Jonsson Comprehensive Cancer Center and BSCRC Ablon Scholars Award, Rose Hills Foundation Stem Cell Innovation Award, the BSCRC and California NanoSystems Institute Stem Cell Nano-Medicine Initiative Planning Award, the BSCRC transformative technology development award to Y. Z.; the National Institute of Neurological Disorders and Stroke (R01NS137919 to JW), National Cancer Institute (R01CA253215 to JW). This work was also supported by the UCLA AIDS Institute, the James B. Pendleton Charitable Trust, and the McCarthy Family Foundation. The graphic figure was created with BioRender.com.

---

**Supplementary Fig. 1.**
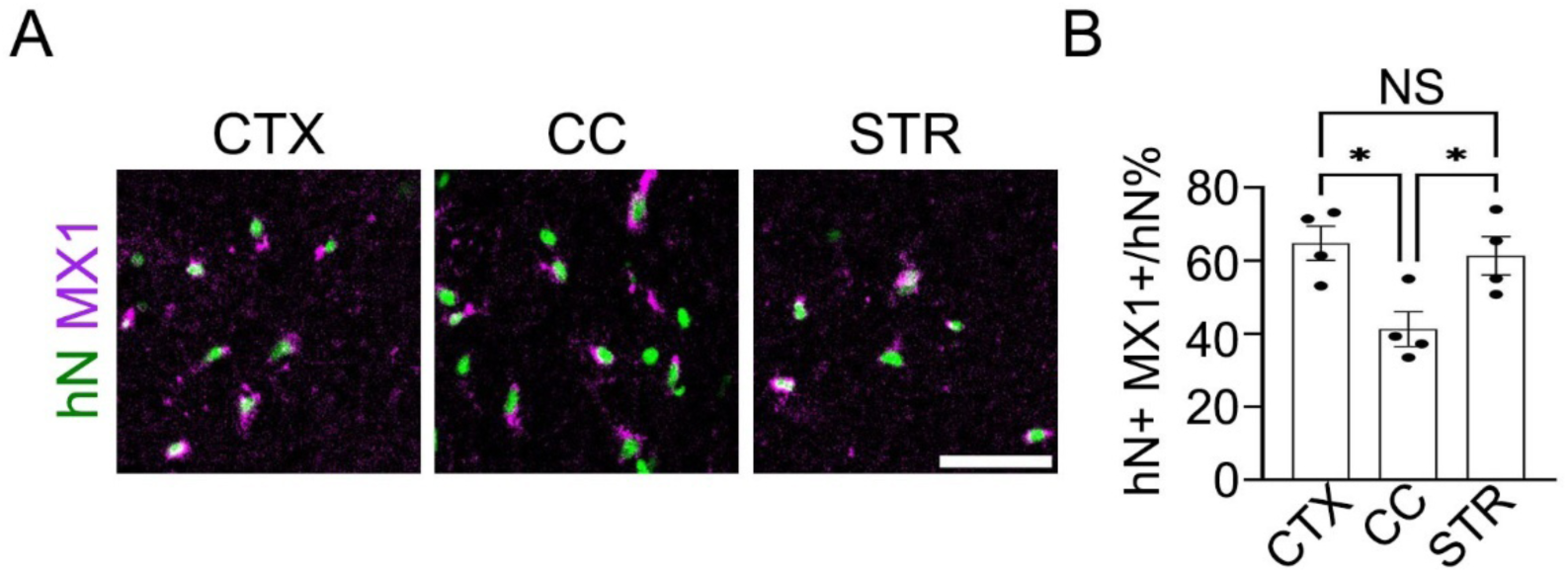
Regional heterogeneity in the adoption of immune oligodendroglia state. (A) Human nuclei (hN) and human MX1 double-staining in cortex (CTX), corpus callosum (CC), and striatum (STR) shows that a smaller proportion of human cells in the CC adopt an immune oligodendroglial state compared to CTX and STR. Scale bar: 50 µm. (B) Quantification of the percentage of hN⁺ cells expressing MX1 in different brain regions (n = 4). One-way ANOVA with Tukey’s multiple comparison correction: CTX vs CC, p = 0.0244; CC vs STR, p = 0.0207; CTX vs STR, p = 0.6147.

**Supplementary Fig. 2.**
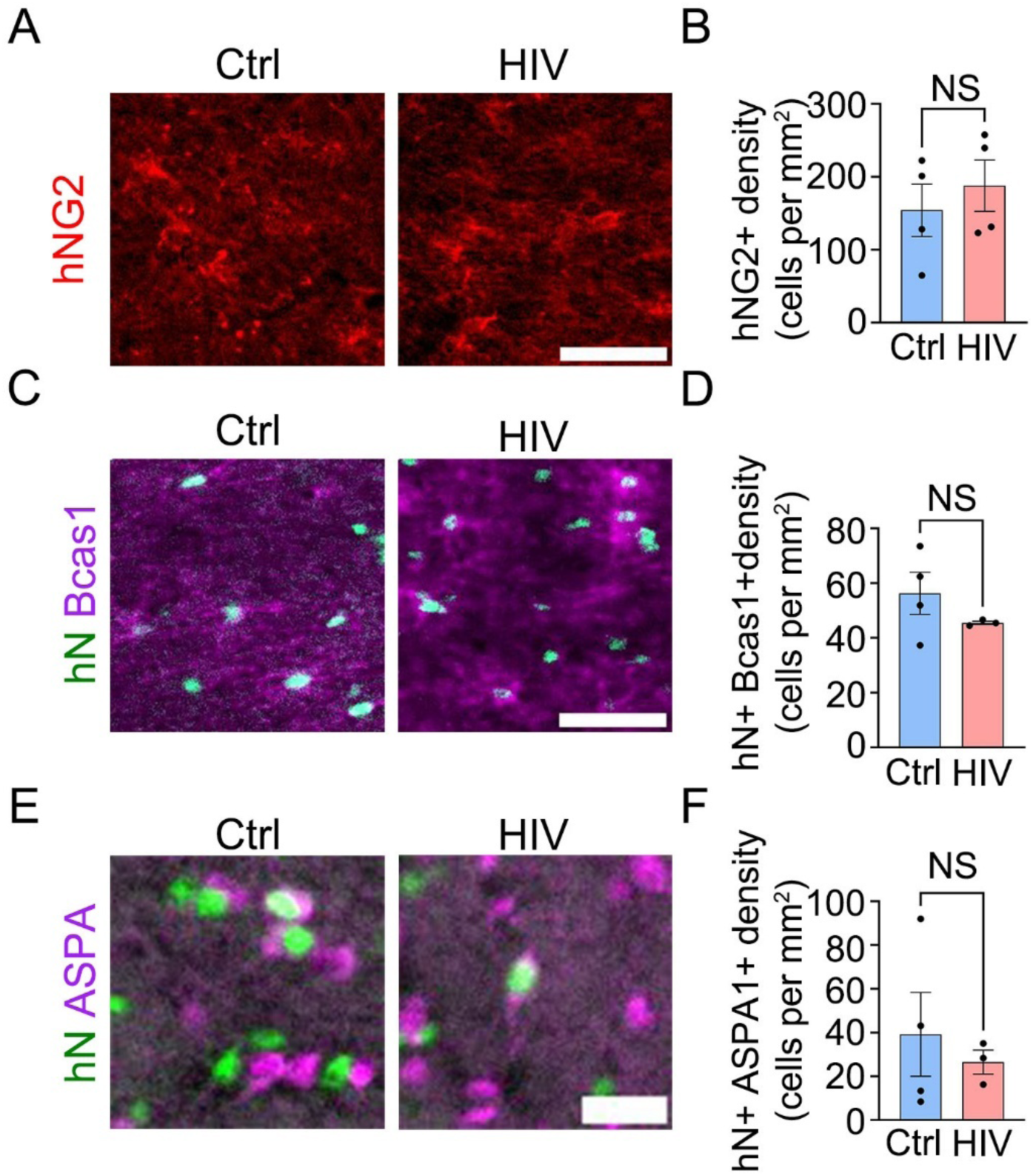
HIV infection does not alter the densities of human oligodendroglia in the chimeric mouse brain. (A) Immunostaining for human NG2 in the corpus callosum shows no change in the densities of human oligodendrocyte precursor cells after HIV infection. Scale bar: 50 µm. (B) Quantification of hNG2⁺ cell density in the corpus callosum (Ctrl: n = 4; HIV: n = 4; unpaired t test, p = 0.5263). (C) Human nuclei (hN) and Bcas1 double-staining shows no change in immature human oligodendrocyte (hN⁺Bcas1⁺) density after HIV infection. Scale bar: 50 µm. (D) Quantification of hN⁺Bcas1⁺ cell density (Ctrl: n = 4; HIV: n = 3; unpaired t test, p = 0.2914). (E) Human nuclei and ASPA double-staining shows no change in mature human oligodendrocyte (hN⁺ASPA⁺) density after HIV infection. Scale bar: 20 µm. (F) Quantification of hN⁺ASPA⁺ cell density (Ctrl: n = 4; HIV: n = 3; unpaired t test, p = 0.6085).

